# Digital ischemia: Etiologies and long-term follow up. A cohort study of 323 patients

**DOI:** 10.1101/468736

**Authors:** Alizée Raimbeau, Marc-Antoine Pistorius, Yann Goueffic, Jérôme Connault, Pierre Plissonneau-Duquene, Blandine Maurel, Jean Reignier, Karim Asehnoune, Mathieu Artifoni, Quentin Didier, Giovanni Gautier, Jean-Nöel Trochu, Bertrand Rozec, Chan N’Gohou, Cécile Durant, Pierre Pottier, Bernard Planchon, Christian Agard, Olivier Espitia

## Introduction

> Upper extremity digital ischemia (UEDI) is a persistent peripheral vascular disease. Diagnosis remains a real challenge to identify and consequently evaluation of etiologies and determination of therapeutic strategies has proved difficult. The incidence of UEDI is about 2 cases per 100,000 persons per year (1,2) with symptomatic ischemia of the upper extremity being encountered less frequently than lower extremity vascular insufficiency.

The main causes generally identified in literature are cardiac or arterial embolism, local thrombosis, systemic autoimmune connective tissue diseases (especially systemic sclerosis), vasculitis or traumatic injury (3).

Past literature shows that only a few studies have focused on the prevalence of etiologies, in patients diagnosed with digital ischemia, with long-term clinical outcomes thus being rarely discussed. Only two retrospective studies about UEDI, with a cohort of over one hundred patients, were reported (2,4,5).

Moreover, there are only few studies published about long-term follow up of UEDI. Three of them were among specific patients : cancer-induced UEDI (4), hypothenar hammer syndrome (6), systemic sclerosis (7)). Only two studies were about general follow up of UEDI, both with a limited number of cases (8,9).

The purpose of this retrospective study was to evaluate etiologies in a large cohort and to assess long-term clinical outcomes of patients diagnosed with digital ischemia.

## Material and Methods

In this retrospective cohort study, we included patients hospitalized for UEDI and outpatients visiting for UEDI in a university hospital between January 1, 2000 and December 31, 2016. We identified patients, based on IDC-10 (I742, T345 and I744 codes) through the medical data processing department. This study was approved by the Local Ethics Review committee. UEDI inclusion criterion was the first episode of UEDI diagnosed, defined by: painful and digital blanching, cyanosis, ulceration of the fingers, gangrene or change of cutaneous temperature, due to vascular pathology. We include all UEDI cases, not only those limited to fingers.

Clinical assessment included demographic data (age, gender, occupation), medical history, cardiovascular risk factors (CVRF), substance abuse, number of recurrence, long-term clinical outcomes and cardiovascular events (stroke, myocardial ischemia, lower limbs ischemia).

Each etiology was determined as follow: Systemic Sclerosis cases (SSc) were diagnosed through ACR/EULAR 2013’s criteria (10); thromboangiitis obliterans cases (TAO) by Mills 2003’s criteria (11); cardio-embolic (CE) diseases were diagnosed through ultrasound criteria and Holter monitoring; vasculitis cases were diagnosed by histology or by cryoglobulinemia positivity or ANCA positivity and further with giant cell arteritis and Takayasu’s Arteritis by ACR or Ishikawa’s criteria; thrombophilias were determined through specific tests; cancer were diagnosed by histology and molecular biology; iatrogenic etiology was defined by intrinsic and extrinsic probability for drug-induced iatrogenic ischemia and by corresponding temporality, for technical-induced iatrogenic ischemia (such as artery cannulation) and finally hypothenar hammer syndrome was identified with the help of occupational health service. When criteria were not found, diagnosis was made following intrinsic and extrinsic causalities. Idiopathic ischemia was defined as such after negative clinical, laboratory and imaging examination. When several etiologies could be incriminated, we chose to include the UEDI case in the etiology most susceptible to be responsible for.

Long-term follow-up was performed by chronological analysis of medical records and by regimenting telephone interviews with patient’s general practitioners.

### Statistical analysis

Results were expressed as mean ± standard deviation (SD) or as median, range. Categorical variable for the groups were compared using chi-square tests or Fisher’s exact tests when any of the expected cell counts of a 2×2 table was less than 5. Comparisons of quantitative variables were performed using Student’s *t-t*est. ANOVA test was performed with a Bonferroni correction after test to individualize group differences. A *p*-value < 0.05 was considered statistically significant. Survival curves were developed to compare cardio-embolic, SSc, idiopathic and TAO groups. Only new events were considered. Survival free of digital ischemia, cardiovascular event (stroke, acute myocardial infarction or acute lower limb arteritis), fingers amputation and global survey were compared. Kaplan-Meier curves were then produced and a log-rank test was performed to analyze the event-free survival. All calculations were done using Graph Pad Prism 6.

## Results

### Patients’ characteristics

323 patients were analyzed. The etiologies of digital ischemia are presented in Table 1

**Table 1:** Distribution of etiology at the end of the follow-up : SSc: systemic sclerosis, TAO: Thromboangiitis Obliterans

| Etiology of digital ischemia | Number of patients (%) |
| --- | --- |
| Cardio embolic | 59 (18.3%) |
| Systemic sclerosis | 52 (16.1 %) |
| Idiopathic | 38 (11.7%) |
| TAO | 30 (9.3%) |
| Iatrogenic | 30 (9.3 %) |
| Cancer | 20 (6.2%) |
| Atherosclerosis | 11 (3.4%) |
| Connective Tissue Disease except SSc | 10 (3.1%) |
| Vasculitis | 10 (3.1%) |
| Traumatic | 9 (2.8%) |
| Thoracic outlet syndrome | 8 (2.5%) |
| Thrombophilia | 7 (2.2%) |
| Infectious disease | 7 (2.2%) |
| Hypothenar hammer syndrome | 6 (1.9%) |
| Vasospasm | 5 (1.5%) |
| Peripheral embolism | 3 (0.9%) |
| Other | 18 (5.6%) |

At the end of the follow-up (average follow-up time: 29.3 months), UEDI associated with cardio embolic disease (DICE) was the highest-occurring etiology in this cohort with 59 patients (18.3%), SSc was the second highest occurring at 16% (52 cases). 38 cases (11.7%) remained undetermined and were classified as idiopathic UEDI. TAO and iatrogenic UEDI were the fourth highest occurring, equally placed in 9.3% of UEDI.

Half of iatrogenic patients were divided up as followed: 30% of cases (9 patients) had vascular steal syndrome from arteriovenous fistula creation (AVF) and 22.2% (6 patients) presented UEDI after vasoconstrictive drug use as norepinephrine in critical care department. Other half of iatrogenic UEDI (15 patients) reported more diffusely : 5 with radial artery puncture (1 after coronary angiography, 1 after renal artery angioplasty, 1 after a blood gas analysis; 1 with invasive arterial central blood pressure; 1 after insertion of a catheter) 3 cases of radiation-induced stenosis; 2 other adverse effects of AVF creation: 1 arterial thrombosis and 1 “tourniquet effect” of a contention put for AVF creation’s hyper flow; 2 cases of other drug-induced side effects (mesalazine and tocilizumab); 1 arterial thrombosis after a local corticosteroid injection of a carpal tunnel syndrome therapy; 1 thrombosis of a carotid-carotid bypass after repeating angio-CT; and 1 case of graft versus host reaction with a systemic sclerosis-like presentation.

UEDI was associated with cancer in 6.2% of cases. Median time between diagnosis of DI and diagnosis of cancer was 14.3 months [1.5-77.1 months]. Malignant hemopathies (45% of all neoplasias) were represented as follows: 30% of myeloproliferative diseases (3 Polycythemia Vera, 1 essential thrombocythemia, 1 secondary myelofibrosis after essential thrombocythemia, 1 myeloid leukemia); 2 multiple myelomas; 1 acute leukemia. Solid tumors (55% of all neoplasias) were distributed as follows: 35% of adenocarcinomas (2 colonic adenocarcinomas, 1 esophageal adenocarcinoma, 1 cholangiocarcinoma, 2 metastatic pulmonary adenocarcinomas, 1 ovarian adenocarcinoma); 2 hepatic neoplasias; 1 Squamous cells carcinoma; and 1 thymoma.

Connective Tissue Disease, without systemic sclerosis, accounted for 3.1% of cases of digital ischemia with 5 cases of mixed connective tissue disease and anti-synthetase syndrome, dermatomyositis, systemic lupus erythematosus, Sjögren’s syndrome, and unclassifiable CTD diagnosed on hepatic biopsy (1 case each).

Among the 11 cases of atherosclerosis’ UEDI, we found 10 cases of atherosclerosis stenosis without subclavian aneurysm and 1 case of aortic arch aneurysm.

UEDI was associated with vasculitis in 6.2% of cases: 3 Giant cells arteritis, 2 Takayasu’s arteritis, 2 cryoglobulinemia, 1 necrotizing small vessel vasculitis and 2 unclassifiable vasculitis. Thrombophilias were represented mostly by antiphospholipid syndrome (6 cases) as well as one case of antithrombin deficiency. Infectious causes were four cases of purpura fulminans and 3 cases of endocarditis.

The 5 cases of vasospasm were 3 cases of narcotic intra-arterial injections (cocaine, heroin) and 2 patients who misused buprenorphine, one with a cold exposure during work-time and one extreme vasospasm on peripheral vascular disease. Peripheral embolisms showed two subclavian artery thrombosis and one aortic arch thrombosis.

Finally, 18 cases were consider as « Other » defined by: 4 cases of calciphylaxis, 1 frost injury, 1 circulatory failure, 1 pre-necrotizing state on severe acrocyanosis, 1 abscess on needle stuck in muscle in a former drug addict, 1 disseminated intravascular coagulation, 2 cases of cold agglutinin disease and 2 cases of severe carpal tunnel syndrome. There were also 3 arteriovenous malformations and 2 congenital stenosis (one brachial artery stenosis, one brachiocephalic trunk stenosis). Traumatic causes were mostly due to violent direct traumas: car accident, stab wounds, attempted suicide, and work accidents.

The main characteristics of the population, as well as characteristics of the four most represented etiologies of UEDI are presented in Table 2.

**Table 2:**
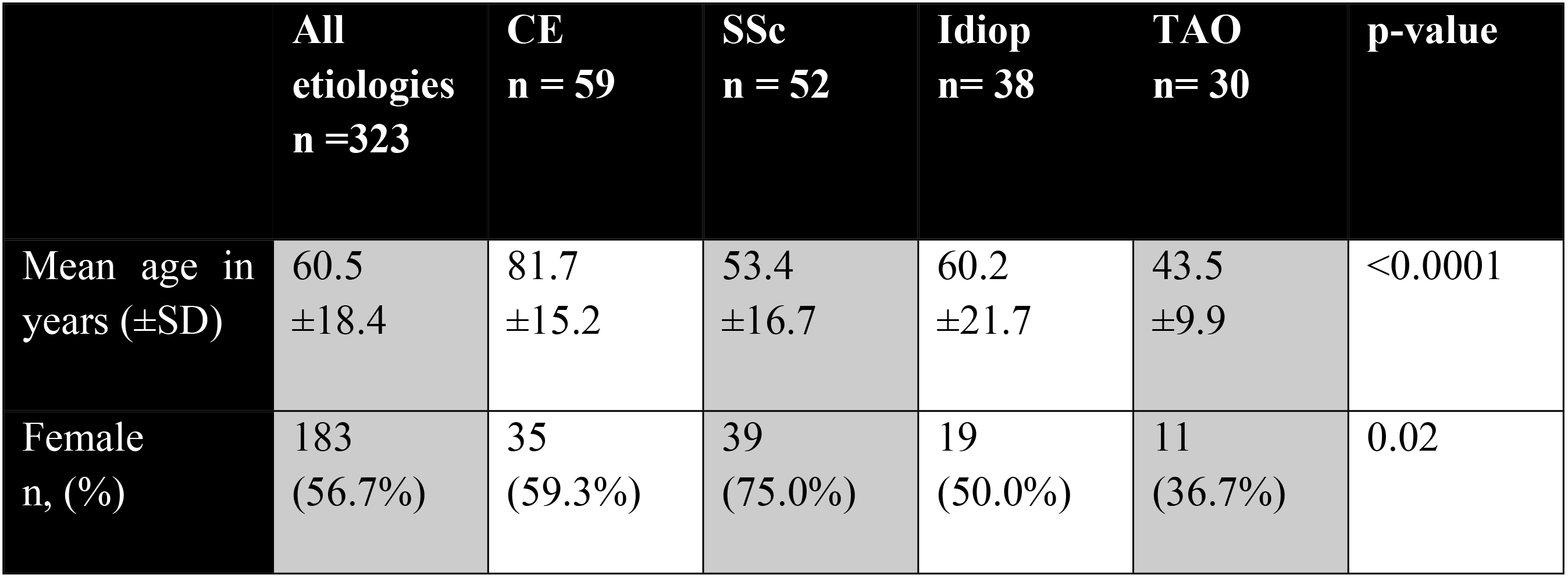
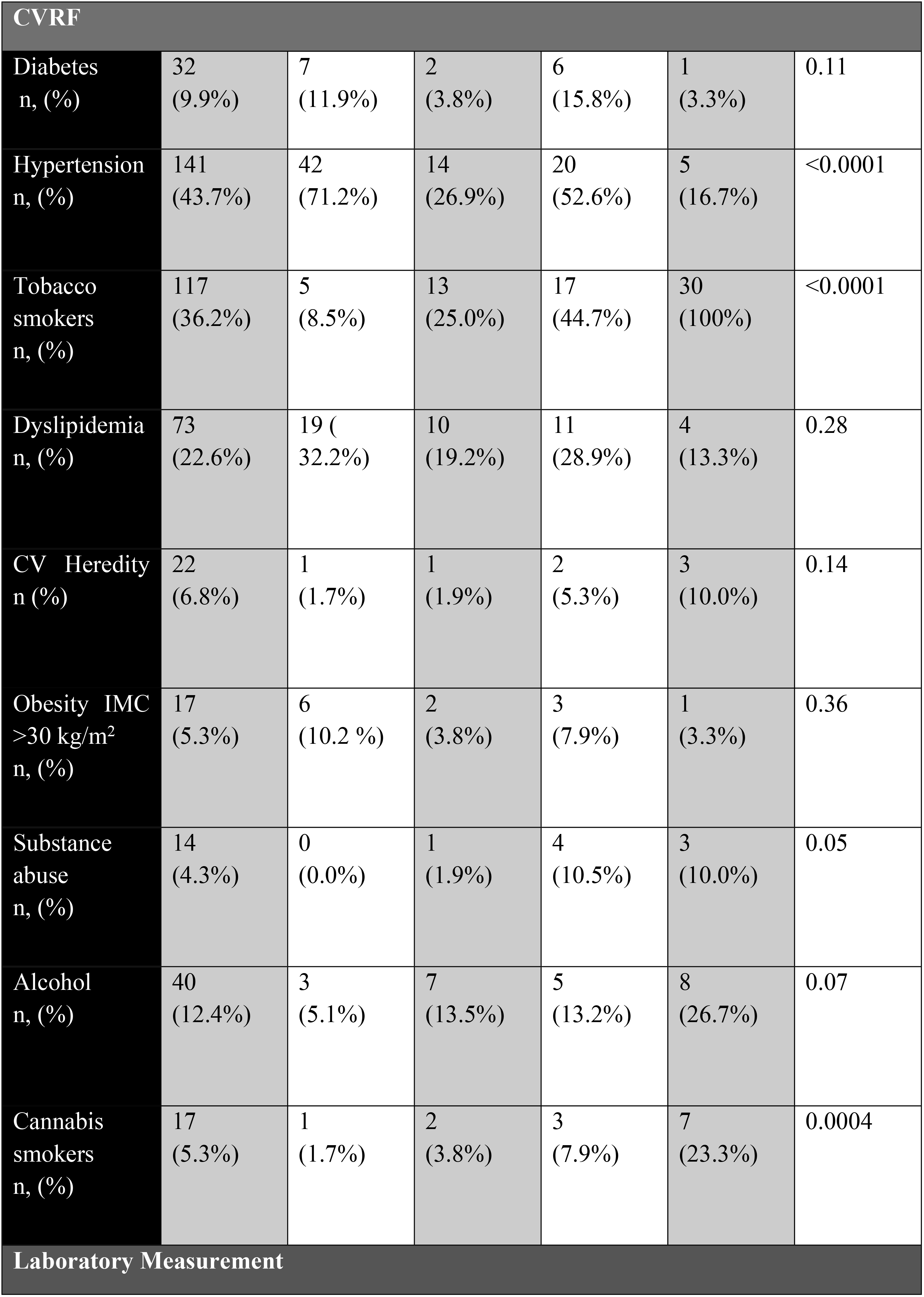
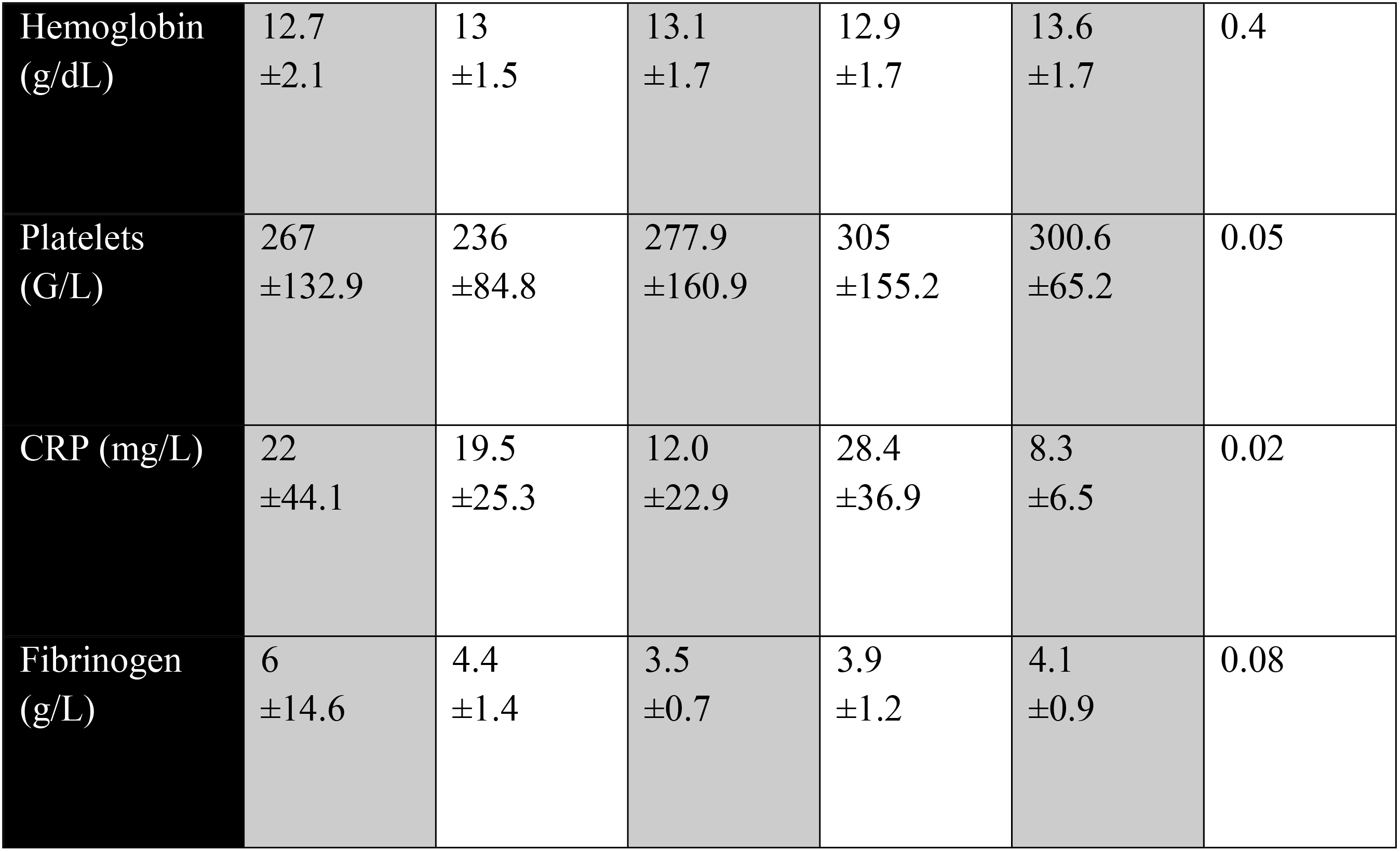
General characteristics of four most commonly occurring etiologies. CVRF: cardiovascular risk factor, CE: Cardio-embolic, Idiop: Idiopathic, SSc: Systemic Sclerosis, TAO: Thromboangiitis obliterans

|  | All etiologies<br>n =323 | CE<br>n = 59 | SSc<br>n = 52 | Idiop<br>n= 38 | TAO<br>n= 30 | p-value |
| --- | --- | --- | --- | --- | --- | --- |
| Mean age in years (±SD) | 60.5<br>±18.4 | 81.7<br>±15.2 | 53.4<br>±16.7 | 60.2<br>±21.7 | 43.5<br>±9.9 | <0.0001 |
| Female<br>n, (%) | 183<br>(56.7%) | 35<br>(59.3%) | 39<br>(75.0%) | 19<br>(50.0%) | 11<br>(36.7%) | 0.02 |

| CVRF |  |  |  |  |  |  |
| --- | --- | --- | --- | --- | --- | --- |
| Diabetes<br>n, (%) | 32<br>(9.9%) | 7<br>(11.9%) | 2<br>(3.8%) | 6<br>(15.8%) | 1<br>(3.3%) | 0.11 |
| Hypertension<br>n, (%) | 141<br>(43.7%) | 42<br>(71.2%) | 14<br>(26.9%) | 20<br>(52.6%) | 5<br>(16.7%) | <0.0001 |
| Tobacco<br>smokers<br>n, (%) | 117<br>(36.2%) | 5<br>(8.5%) | 13<br>(25.0%) | 17<br>(44.7%) | 30<br>(100%) | <0.0001 |
| Dyslipidemia<br>n, (%) | 73<br>(22.6%) | 19 (32.2%) | 10<br>(19.2%) | 11<br>(28.9%) | 4<br>(13.3%) | 0.28 |
| CV Heredity<br>n (%) | 22<br>(6.8%) | 1<br>(1.7%) | 1<br>(1.9%) | 2<br>(5.3%) | 3<br>(10.0%) | 0.14 |
| Obesity IMC<br>>30 kg/m<sup>2</sup><br>n, (%) | 17<br>(5.3%) | 6<br>(10.2 %) | 2<br>(3.8%) | 3<br>(7.9%) | 1<br>(3.3%) | 0.36 |
| Substance<br>abuse<br>n, (%) | 14<br>(4.3%) | 0<br>(0.0%) | 1<br>(1.9%) | 4<br>(10.5%) | 3<br>(10.0%) | 0.05 |
| Alcohol<br>n, (%) | 40<br>(12.4%) | 3<br>(5.1%) | 7<br>(13.5%) | 5<br>(13.2%) | 8<br>(26.7%) | 0.07 |
| Cannabis<br>smokers<br>n, (%) | 17<br>(5.3%) | 1<br>(1.7%) | 2<br>(3.8%) | 3<br>(7.9%) | 7<br>(23.3%) | 0.0004 |
| Laboratory Measurement |  |  |  |  |  |  |
| Hemoglobin (g/dL) | 12.7<br>±2.1 | 13<br>±1.5 | 13.1<br>±1.7 | 12.9<br>±1.7 | 13.6<br>±1.7 | 0.4 |
| Platelets (G/L) | 267<br>±132.9 | 236<br>±84.8 | 277.9<br>±160.9 | 305<br>±155.2 | 300.6<br>±65.2 | 0.05 |
| CRP (mg/L) | 22<br>±44.1 | 19.5<br>±25.3 | 12.0<br>±22.9 | 28.4<br>±36.9 | 8.3<br>±6.5 | 0.02 |
| Fibrinogen (g/L) | 6<br>±14.6 | 4.4<br>±1.4 | 3.5<br>±0.7 | 3.9<br>±1.2 | 4.1<br>±0.9 | 0.08 |

Mean age was 60.5 year old; (14-102 years old); One third were active smokers (36.2%). Highest occurring CVRF was hypertension with 43.7%, then dyslipidemia (22.6%) and diabetes (9.9%).

DICE patients were significantly older than all others groups (p<0.001) except the neoplasia group. They presented significantly more hypertension than SSc and TAO groups (p<0.0001). The DICE group comprised also fewer tobacco smokers than in the idiopathic group (p<0.0001). In the SSc group, there were significantly more women than in the TAO group (p=0.0006), idiopathic group (p=0.03) and iatrogenic group (p=0.04). TAO patients were significantly younger than those in the iatrogenic group (p<0.001) and in the neoplasia group (p<0.01). There were also significantly more tobacco smokers than in all other groups (p<0.0001) and more cannabis smokers than in all other groups except idiopathic (DICE (p=0.002), neoplasia (p=0.03), SSC and iatrogenic (p=0.01)). In TAO group, there were significantly more men than in the DICE (p=0.048) and neoplasia (p=0.04) groups and patients were also significantly less hypertensive than in all other groups except SSc group (p=0.42).

### Surgical revascularization according to etiology

**Figure 1:**
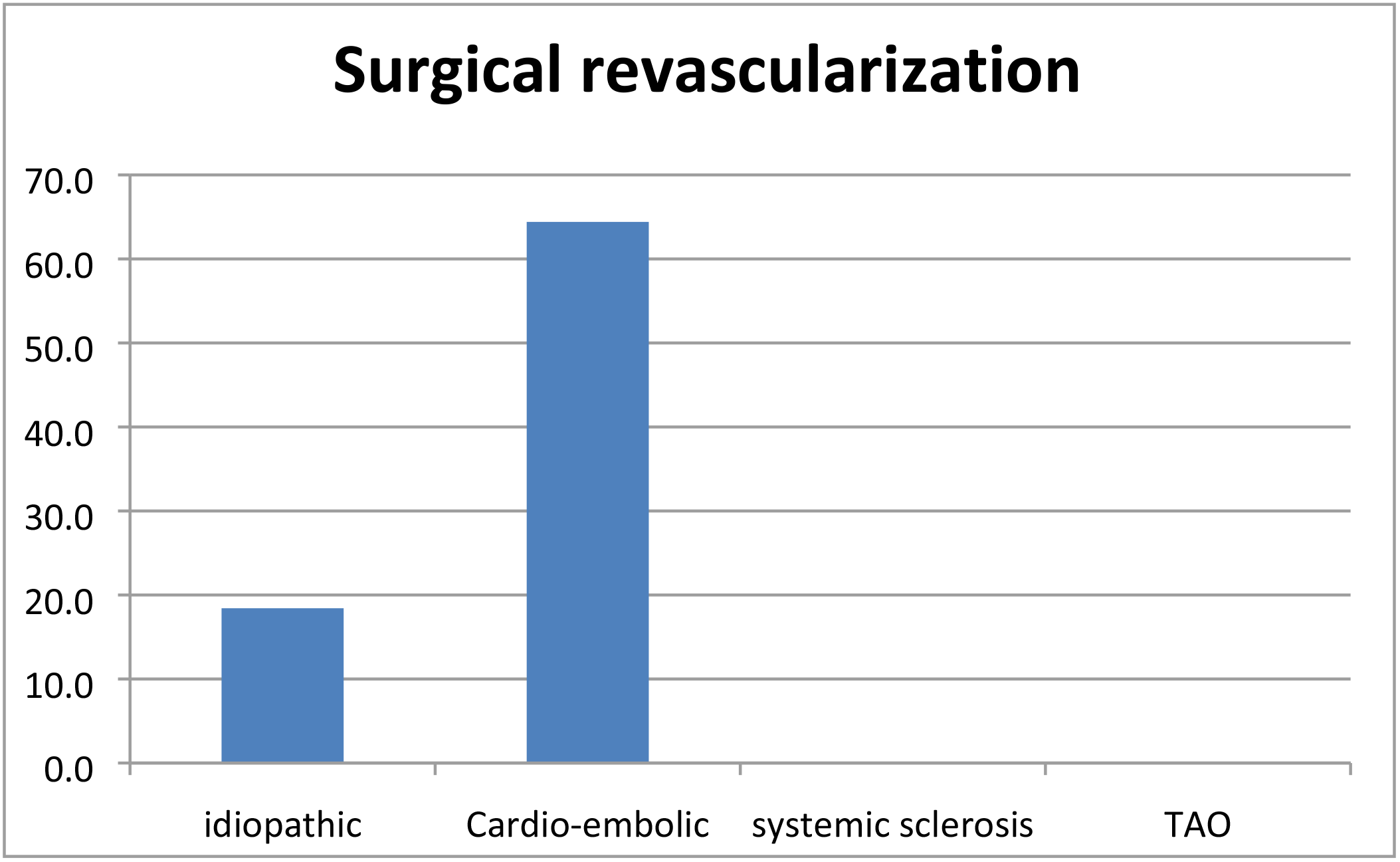
Surgical revascularization according to etiology. TAO : Thromboangiitis Obliterans.

Successful surgical revascularizations were significantly more often than all other groups (p<0.0001).

Idiopathic group had also significantly more surgical revascularization than SSc group (p=0.002) and TAO group (p =0.02).

### Recurrences, digital amputations, cardiovascular morbidity and mortality

The rate of recurrence was various and etiology-dependent (Figure 2A). Recurrence of UEDI was significantly greater in SSc group (n= 7) than in all other groups (p <0.0001 vs idiopathic and DICE, p=0.0004 vs iatrogenic p=0.003 vs TAO and p=0.007 vs neoplasia). In the TAO group, recurrence was significantly greater (n=7) than in DICE (n=1) (p=0.005). There were only 3 recurrences in idiopathic group.

**Figure 2:**
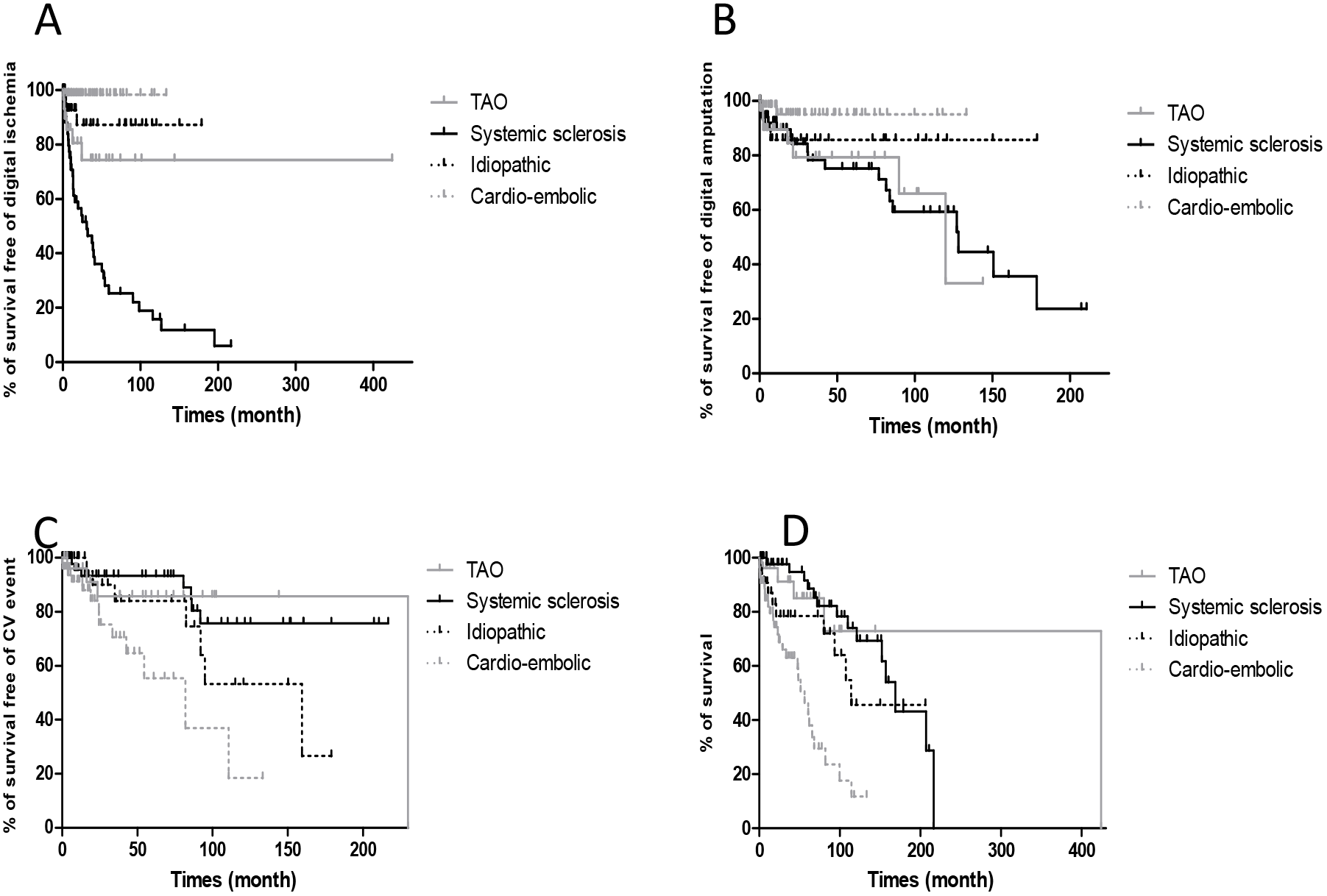
Long-term follow up and survival curves **A**. Survival free of recurrence of UEDI; **B**. Survival free of amputations and auto-amputations; **C.** Survival free of CV events, **D**. Global survey. CV: Cardiovascular, UEDI: Upper Extremity Digital ischemia, TAO: Thromboangiitis Obliterans.

Frequencies of digital amputations and auto-amputations varied, depending on etiologies (Figure 2B). The cumulated rate of digital amputation and auto-amputation was higher in SSc group (n=18) (p=0.02) and TAO (n=7) (p=0.03) than in DICE group (n=2). In terms of survival (free of amputation) features, there were no significant differences between the other groups. There were only 4 amputations and auto-amputations in the idiopathic group.

CV events also varied (Figure 2C) : the DICE group had more CV events (n= 17) than idiopathic UEDI (n=8) (p=0.04), SSc (n=7) (p=0.0005) and TAO (n=4) (p=0.03).

Global survival was different depending on etiology (Figure 2D). Mortality was more frequent in DICE (n=27) than SSC (n=14) (p<0.0001), idiopathic (n= 10) (p=0.0093) and TAO (n=5) (p=0.0007) groups. However, there was no difference between idiopathic group and: SSc (p=0.12), iatrogenic (p=0.51) and TAO group (p = 0.29).

## Discussion

This study on digital ischemia is the largest cohort ever reported with 323 patients encountered in tertiary center recruitment in vascular pathology.

At the end of the follow-up DICE was the highest-occurring etiology in this cohort, the second highest occurring was SSc. Idiopathic UEDI were the third highest occurring, TAO and iatrogenic UEDI were the fourth highest occurring, equally placed.

DICE recurs only in a small number of cases in this study. Conversely, in this study, patients with SSC and TAO presented higher number of recurrence. Similarly, patients with SSC/TAO-associated UEDI presented with higher number of cases requiring amputation than DICE patients. This phenomenon may be due to the chronicity of vascular lesions and the disappearance of microcirculatory reserves, associated with distal involvement. In contrast, DICE occurs in arteries of larger caliber, usually without pre-existing lesions, which underpins the success of surgical management and the low amputation rate. DICE, however, is likely to be a marker of overall cardiovascular risk as patients with DICE have more cardiovascular events and have higher mortality than other UEDI groups.

This study showed proximity between DICE and idiopathic UEDI groups. Indeed, there are no survival differences without recurrence of UEDI and no difference in amputation rates between DICE and idiopathic UEDI groups. However, the DICE group had higher mortality and cardiovascular events whilst idiopathic UEDI tended to showcase more cardiovascular events than other groups of UEDI which might suggest that idiopathic UEDI may well be unidentified DICE although in this study, we did not find a significant difference. The low rate of recidivism is also in favor of this proximity.

The etiologies of the digital ischemia are numerous and, according to previous studies, there are differences in the distribution of etiologies (Table 3). Our results demonstrate that 18.3% of digital ischemia was caused by cardio-embolic disease, 16% were due to systemic sclerosis and 11.7%, 9.3% and 9.3% of idiopathic, TAO and iatrogenic origins respectively. Table 3 shows the distribution of digital ischemia etiologies in the largest series of literature ever reported.

**Table 3:** Comparison of etiologies in other studies (CTD: Connective Tissue Disease, CE: Cardioembolic HHS: Hypothenar hammer syndrome, Idiop: Idiopathic, Iatro: iatrogenic, NA: Not Available, Néo: neoplasia, SSC: Systemic sclerosis, TAO: Thromboangiitis obliterans.

|  | All<br>CTD | CE | SSc | Idiop | Iatro | TAO | Cancer | Atherosclerosis | vasculitis | HHS |
| --- | --- | --- | --- | --- | --- | --- | --- | --- | --- | --- |
| **Present study**<br>**n=323** | 19.2% | 18.3% | 16.0% | 11.7% | 9.3% | 9.3% | 6.2% | 3.4% | 3.1% | 1.6% |
| **Carpentier and al 2005**<br>**(2,5)**<br>**n = 278** | 33.0% | 8.6% | 26.0% | 4.0% | 0.3% | 10.0% | 4.0% | 15.0% | 0.0% | 15.0% |
| **Le Besnerais and al. 2014**<br>(4)<br>**n= 100** | 10.0% | 8.0% | 10.0% | 10.0% | NA | 22.0% | 15.0% | 4.0% | 0.0% | 13.0% |
| **Cailleux and al. 1999**<br>(12)<br>**n = 96** | 26.0% | 6.2% | 26.0% | 10.5% | NA | 11.5% | 5.0% | NA | NA | NA |
| **Vayssairat and al. 1977**<br>(13)<br>**n=86** | 48.8% | NA | 39.0% | 19.0% | NA | 4.6% | 3.4% | 8.0% | 8.1% | 2.3% |
| **Cailleux and al. 1994**<br>(14)<br>**n= 45** | 27.0% | 9.0% | 0.0% | 4.0% | 0.0% | 13.0% | 2.0% | 16.0% | 0.0% | 11.0% |
| **Keo and al. 2011** (9)<br>**n = 36** | 42.0% | 6.7% | 14.0% | 22.0% | 2.8% | 0.0% | 0.0% | 11.0% | 0.0% | 33.0% |
| **Abdallah and al. 2009**<br>(15)<br>**n=25** | 36.0% | 4.0% | 12.0% | 4.0% | 0.0% | 20.0% | 8.0% | 20.0% | 12.0% | 0.0% |

This study found CE disease to be the highest-occurring etiology of UEDI accounting for 18.3% of cases, while in literature, this etiology distribution rate varied between 4 and 9%. Previous studies have also shown connective tissue diseases to be the highest occurring etiology of UEDI (2,4,9,12–15). In this study, however, all CTD accounted for 19.2% of UEDI whilst systemic sclerosis accounted for 16% of cases. Indeed most of the data are derived from studies of digital ischemia, concerning a specific etiology or a range of clinical outcomes from the internal medicine department, where connective tissue disorders are very strongly represented. Moreover, our inclusion criteria included hand ischemia rather than limited finger ischemia, as typically employed, which may have contributed to a greater recruiting of CE patients. Our findings showed that UEDI of undetermined origin, after well-conducted explorations, is not uncommon and accounts for 11.7% in our study; in the literature, idiopathic digital ischemia accounts for 4-22% of cases.

Iatrogenic cause was the fourth highest-occurring etiology, in this study, with a rate of 9.6% vs 0 to 2.8% in literature (2,4,9,12–15); where for 3 of 7 other studies, iatrogenic etiology was not a research focus. The stronger representation, in this study, is linked to a broad recruitment of patients, in critical care units, with more drug-induced iatrogenic UEDI. Additionally, the results of previous studies may be limited by the era in which research was carried out as intra-arterial proceedings have developed significantly, particularly in recent years.

Cancer-associated UEDI was found in 2 to 15% of cases. In this study, as in Le Besnerais et al (4), the most common histological type was adenocarcinoma. Myeloproliferative diseases were also frequent here, accounting for 30% of digital ischemia associated with cancer. The time between UEDI and cancer diagnosis varied, with it taking 2 months in a study with intensive initial investigation (4) and 14.3 months, in this study, after a clinical examination, standard blood test and clinical follow-up.

### Limits

In this study, patients were included retrospectively in a large time-lapse of sixteen years and so there was data loss. We presented episodes of UEDI from tertiary center recruitment in vascular pathology, where prevalence of CTD and multiple comorbidities are higher. Because diagnosis of occupational pathologies is carried out upstream of our center, they are under-represented in our study while they represent up to 33% in some UEDI studies.

## Conclusion

This study describes the largest cohort of UEDI ever reported. For the first time, this study showed that cardio-embolic diseases are the highest occurring of the UEDI etiologies studied. This study also found that iatrogenic cause of UEDI was frequent and increased with the use of vasopressor drugs and multiplication of endoluminal arterial procedures. The long-term follow-up showed: UEDI associated with SSc had a poor local prognosis and DICE presented poor general prognosis. UEDI with SSc and TAO presented with more recurrence. When idiopathic UEDI shows recurrence, those etiologies have to be researched as a priority. This research is even more important because of the higher risk of clinical course towards amputations in those etiologies.

## Abbreviations

UEDI: Upper Extremity Digital Ischemia
DICE: Digital ischemia due to Cardio-Embolic disease
SSc: Systemic Sclerosis
TAO: ThromboAngiitis Obliterans

## References

1. Bae M, Chung SW, Lee CW, Choi J, Song S, Kim S. Upper Limb Ischemia: Clinical Experiences of Acute and Chronic Upper Limb Ischemia in a Single Center. Korean J Thorac Cardiovasc Surg. août 2015;48(4):246–51.

2. Carpentier PH, Guilmot JL, Hatron PY, Levesque H, Planchon B, Vayssairat M, et al. Nécroses et artériopathies digitales. J Mal Vasc. sept 2005;30(4):29–37.

3. Senet P. Diagnostic des acrosyndromes vasculaires. Ann Dermatol Vénéréologie. août 2015;142(8–9):513–8.

4. Le Besnerais M, Miranda S, Cailleux N, Girszyn N, Marie I, Lévesque H, et al. Digital Ischemia Associated With Cancer. Medicine (Baltimore). 22 août 2014;93(10).

5. Delcey V, Michon-Pasturel U, Cailleux N, Hatron PY, Hachulla E, Devulder B, et al. Nécroses et ischémies digitales du membre supérieur :étude rétrospective de 278 observations. Rev Médecine Interne. 1 déc 2001;22(Supplement 4):445s–446s.

6. Marie I, Hervé F, Primard E, Cailleux N, Levesque H. Long-term follow-up of hypothenar hammer syndrome: a series of 47 patients. Medicine (Baltimore). nov 2007;86(6):334–43.

7. Tolosa-Vilella C, Morera-Morales ML, Simeón-Aznar CP, Marí-Alfonso B, Colunga-Arguelles D, Callejas_Rubio JL, et al. Digital ulcers and cutaneous subsets of systemic sclerosis: Clinical, immunological, nailfold capillaroscopy, and survival differences in the Spanish RESCLE Registry. Semin Arthritis Rheum. 1 oct 2016;46(2):200–8.

8. Marchal A, Mahé E, Sin C, Bagan P, Bilan P, Linder J-F, et al. Acute finger ischemia: A retrospective study of 13 patients. Ann Dermatol Vénéréologie. mai 2015;142(5):332–9.

9. Keo HH, Umer M, Baumgartner I, Willenberg T, Gretener SB. Long-term clinical outcomes in patients diagnosed with severe digital ischemia. Swiss Med Wkly. 18 févr 2011;141:w13159.

10. Sáez-Comet L, Simeón-Aznar CP, Pérez-Conesa M, Vallejo-Rodríguez C, Tolosa-Vilella C, Iniesta-Arandia N, et al. Applying the ACR/EULAR Systemic Sclerosis Classification Criteria to the Spanish Scleroderma Registry Cohort. J Rheumatol. déc 2015;42(12):2327–31.

11. Mills JL. Buerger’s disease in the 21st century: diagnosis, clinical features, and therapy. Semin Vasc Surg. 1 sept 2003;16(3):179–89.

12. Cailleux N, Marie I, Lecomte F, Peillon C, Lévesque H, Courtois H. L’ischémie digitale: une affaire d’internistes. À propos de 96 observations. Rev Médecine Interne. juin 1999;20:s76.

13. Vayssairat M, Fiessinger JN, Housset E. Les nécroses digitales du membre supérieur : 86 cas. Press Med. 1977;(6):931–4.

14. Cailleux N., Levesque H., Gilbert P., Joly P., Lauret Ph., Testart J., et al. Les nécroses digitales du membre supérieur en dehors de la sclérodermie. Etude rétrospective à propos de 45 cas. J Mal Vasc. 1994;19:22-6.

15. Abdallah M, Hamzaoui S, Larbi T, Bouslama K, Harmel A, Ennafaa M, et al. Profil étiologique des nécroses digitales des membres supérieurs : analyse de 25 observations. J Mal Vasc. févr 2010;35(1):12–6.

